# Time series analysis of trial-to-trial variability of MEG power spectrum during rest state, unattented listening and frequency-modulated tones classification

**DOI:** 10.1101/2021.03.15.435429

**Authors:** Lech Kipiński, Wojciech Kordecki

## Abstract

The nonstationarity of EEG/MEG signals is important for understanding the functioning of human brain. From the previous research we know that even very short, i.e. 250—500ms MEG signals are variance-nonstationary. The covariance of stochastic process is mathematically associated with its spectral density, therefore we investigate how the spectrum of such nonstationary signals varies in time.

We analyze the data from 148-channel MEG, that represent rest state, unattented listening and frequency-modulated tones classification. We transform short-time MEG signals to the frequency domain using the FFT algorithm and for the dominant frequencies 8—12 Hz we prepare the time series representing their trial-to-trial variability. Then, we test them for level- and trend-stationarity, unit root, heteroscedasticity and gaussianity and based on their properties we propose the ARMA-modelling for their description.

The analyzed time series have the weakly stationary properties independently of the functional state of brain and localization. Only their small percentage, mostly related to the cognitive task, still presents nonstationarity. The obtained mathematical models show that the spectral density of analyzed signals depends on only 2—3 previous trials.

The presented method has limitations related to FFT resolution and univariate models, but it is not computationally complicated and allows to obtain a low-complex stochastic models of the EEG/MEG spectrum variability.

Although the physiological short-time MEG signals are in principle nonstationary in time domain, its power spectrum at the dominant frequencies varies as weakly stationary stochastic process. Described technique has the possible applications in prediction of the EEG/MEG spectral properties in theoretical and clinical neuroscience.

## 1 Introduction

Time series modeling plays an important role in the description of complex biological systems and has numerous scientific applications. With regard to the electro-(EEG) or magneto-(MEG) encephalography, proposing an adequate model of neural activity, especially in the context of the brain’s response to stimulation with external stimuli, is crucial for understanding the functioning of neuronal connections in the brain, the phenomenon of perception, cognitive functions and is applicable in numerous fields of neuroscience and medicine. This includes areas such as source modeling, seizure detection prediction, memory and attention research, brain-computer interfaces, and a large number of clinical observations in neurology, psychiatry and other medical disciplines.

Many different methods are used in the literature to describe EEG/EMG signals. Traditional evaluation of spontaneous (rest state) brain activity is based on a description of the waveform morphology in the time domain, while in the frequency domain methods based on the Fourier transform are usually used. In turn, the clinically accepted technique for obtaining event-related responses is the averaging method. A significant methodological limitation in such studies is the fact that EEG/MEG signals are nonlinear and non-stationary, i.e. their properties change over time [27, 31]. Therefore, more mathematically advanced methods of analysis had to find their place in neuroscience [32]. Plenty of them are time-frequency analysis, with the use of tools such as wavelets or matching pursuit [7, 8, 17, 22, 25, 47, 49]. Some of them are fractal or entropy based [45, 51]. Others use the independent component analysis and similar techniques [26, 3, 44, 23]. Also, many algorithms based on statistical methods were developed, mostly devoted to the estimation of evoked responses [16, 50, 19, 29, 33, 52]. Among the latter, attention should be paid to the stochastic time series models, dedicated to the univariate [38] or multivariate [18, 37] description of EEG/MEG data. These methods can be applied, among others for predicting changes in signal parameters over time [48].

Moreover, it is not entirely clear how the human brain processes external information. Traditional view on brain evoked-responses generation assumes the so-called an additive model, in which stimulus-induced activity is hidden in the noise associated with the spontaneous EEG, potentials resulting from the processing of accompanying stimuli, including cognitive processes and artifacts [33]. More recent approach is based on stimulus-related changes in the ongoing EEG/MEG oscillations, then the evoked responses reflect rather the synchronization of such oscillations during stimuli processing in neural network [6, 41, 42, 57]. It is worth noting that the data description method is usually selected to match the previously assumed functional model, what may bias the obtained results [56].

So a few questions arise. Firstly, is it not worth looking for methods of analysis that are not burdened with such assumptions? Second, is it possible to better understand the nature of EEG/MEG signals so that new methods of analyzing them are based on their real properties?

In this paper, we have devoted attention to examining some aspects of the variability of MEG activity. For this purpose we applied statistical test to infer about stochastic properties of encephalographic data, similarly as in work [31]. As mentioned above, the bioelectrical activity of the brain is considered a nonstationary process as it is well known that they are strongly time-varying, especially their spectral power. Therefore, we analyzed the trial-to-trial spectrum variability, obtained by a way similar to [28, 39] and finally proposed a wide class of stochastic time series models for its description.

## 2 Aim

Even very short (0.25-0.50 sec) fragments of MEG signals are proved to be nonstationary due to the time-dependent variance [31]. However the covariance matrix is mathematically associated with signal energy and the spectral density, so variance-nonstationarity is equivalent to the time-variability of spectral power of MEG signals and it is wide described in the bibliography for both, spontaneous and event-related EEG/MEG data and have important neurophysiologic consequences. Since the changes in the spectrum are responsible for the nonstationarity of these signals our goal was to find out how the spectral structure of short-term MEG fragments changes. We chose MEG data recorded at rest state, during passive listening to sounds and related to the recognition and classification of frequency-modulated tones so that our calculations could be used to compare whether the indicated spectrum variability is related to auditory information processing and, if, at what level.

Presented studies are the continuation of research published in [31]. Like in the Section 4.3 of cited work, the modern tests for (non-)stationarity were applied to infer about time-varying statistical properties of MEG power spectra. Moreover, the same real data set was used in both analysis. Yet, in opposition to [31] where only 6 channels were taken into account (4 leads above the auditory cortices of both hemispheres, and 2 – frontal and occipital sensors – used as “reference” channels) here the multichannel analysis is performed. The second new approach in the present work is the estimation of stochastic time series models for series of the trial-to-trial variability of spectra power in selected Fourier frequencies. The statistic modeling is used for parametric description of analyzed MEG variability.

## 3 MEG experiment

The MEG experiment was performed at the Special Lab for Non-Invasive Brain Imaging in Leibniz Institute for Neurobiology in Magdeburg, Germany. A whole-head MEG device (Magnes 2500 WH, 4-D Neuroimaging, San Diego, USA) containing 148 magnetometer sensors coupled to DC SQUID’s was used to acquire the data. The measured signals are corrected for environmental noise by means of a weighted subtraction of reference signals detected by additional sensors located in close proximity to the field detectors. In order to avoid any noise contribution from the liquid helium, all pick-up and reference detectors, as well as the 148 SQUID’s, are located in vacuum. The MEG apparatus is located in a magnetically shielded room. Data were sampled at 1017.25 Hz and bandpass filtered during acquisition between 0.1Hz and 100 Hz.

The experiment was intended to acquire MEG signals representing various functional states of the human brain. For this purpose, it was decided to measure: 1. spontaneous brain activity; 2. signals registered during the repeated tonal stimulation (evoked fields); 3. MEG waveforms containing cognitive event-related components connected with perception of frequency-modulated (FM) tones (with their classification task during one session). In each FM sweep the instantaneous frequency changes linearly in time, either upward or downward. A set of 32 FM tones was used, 16 for both categories, presented in random order. The initial frequencies for the rising FMs varied from 0.5 kHz to 2 kHz in steps of 100 Hz, those for falling FMs from 4 kHz to 1 kHz in steps of 200 Hz. Each FM sweep had duration of 500 ms including 10 ms linear ramps both at the beginning and the end.

The acoustic stimulation with FM tones was chosen because they have fundamental role in the human speech as well as in the animal vocalizations. Studies using FM tones provide a possibility to gain insight into the processing of prosodies independent of any speech-specific mechanisms. The FM tones can be easily separated into two natural categories with respect to the underlying direction of frequency modulation, upward or downward. Thus, they also serve as an ideal tool for comparing the auditory cortex response to a passive condition, that is, mere listening to a sequence of FM tones, with an active condition (cognitive task), that is, the directional categorization of exactly the same FM tones. Different activation patterns for the two conditions provide insight into the impact of cognitive aspects (related to FM tone discrimination) on the processing of FM tones in the auditory cortex [57]. The FM tones find their importance not only in MEG studies on auditory pathway yet also in fMRI-based neuroscience [53, 4, 21]. More details about the FM stimuli are given in [34].

There were two stages of the experiment when the FM tones were used. In the first one, the exposure condition (FM/E), the subject had to listen passively to the sequence of FM tones. In the second one, the task condition (FM/T), he was instructed to discriminate the direction of frequency modulation of the same FM tones and to categorize them by pressing one of two response keys. The subject was instructed to signal rising FM sweeps with his left index finger, and falling FM sweeps with his right index finger as soon as he had identified the direction of frequency modulation. Within an experimental session, the presentation of the FM tones was binaural and randomized.

The repeated 1 kHz stimuli had 500 ms duration and 10 ms ramps at the beginning and at the end. The MEG data collected for an acoustic stimulation with such a simple reference stimulus reflect the basic processes behind the stimulus perception in the human brain because the associative processes of stimulus recognition and its classification are intensive only in the very first phase of the examination. Thus, the registered signals are representative for the auditory-cortex processing.

Two sessions in which the spontaneous activity was registered (i.e. during the period when the subject was not exposed to any stimulation) were introduced to serve as the natural reference level for the sessions involving stimulation.

Within each session involving stimuli, the stimulus was presented 96 times. For the 1 kHz stimulation each stimulus was exactly the same, whereas the FM tones were presented randomly, each sweep 3 times, resulting in total number of 96 stimuli (32 × 3). The measurement was continuous, thus every single trial (epoch) corresponding to a single stimulus presentation was equivalent to 2 seconds of registration, where t = 0 corresponds to the stimulus onset for each trial. The spontaneous brain activity was also measured continuously over the period of time, whose length equaled 96 × 2s. Therefore, each experimental session lasted 3 minutes and 4 seconds. Timing diagram of a single trial is given in Fig. 1.

**Figure 1:**
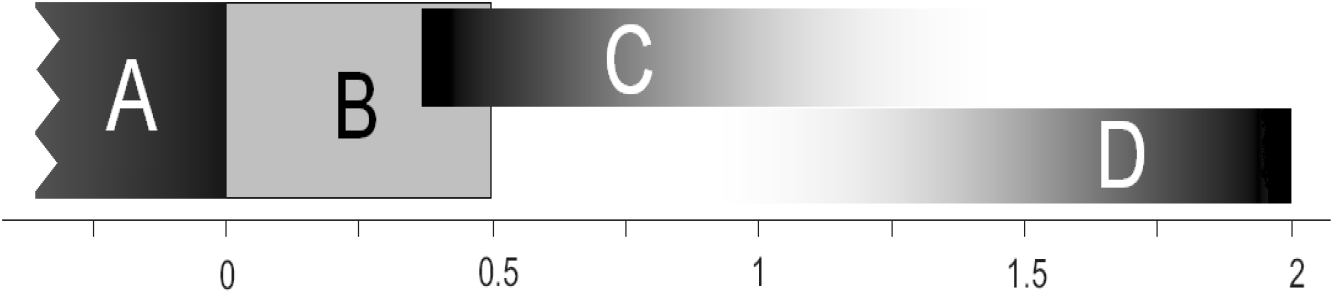
Timing of a single trial: A — final part of a previous trial; B — the period of stimulus presentation; C — response time for classification in the task condition FM/T (absent in the FM/E condition); D — poststimulus period for recovery of the MEG signal. B is absent in session without stimulation. In the FM/T session, C may occasionally overlap with D. C overlaps slightly with the period of stimulus presentation B, as the subject responded already before the end of the stimulus in some trials. Time is given in seconds.

During experiment the acoustic stimuli were generated outside the MEG chamber and delivered to the subject’s ears via two approximately 6-m long plastic tubes. These tubes ended with special ear-molds which were individually chosen to adapt to each subject’s pinnae. The time delay caused by the signal conduction along the plastic tubes was measured as 20ms, and was taken into account in the data analysis. At the beginning of each measurement, the sound pressure level of the acoustic stimulus was adjusted to 90 dB SPL using a 1 kHz sinusoidal tone as reference signal and the subject rated this sound intensity as comfortable.

As the result of measurements from a healthy volunteer, multichannel MEG signals of high quality were obtained for all sessions. All were evaluated by neurologist experienced in description of that kind of data as physiological: the spontaneous MEG activity is typical for resting state in humans and free of any seizure phenomena or sleep waveforms; the evoked fields are typical early cortical auditory responses with proper amplitudes and latencies of main components after averaging, as well as their over-head topography; the event-related fields contain proper cognitive long-latency waveforms after averaging and their topography over head is physiological too. Moreover, the results obtained for the sessions related to acoustic stimulation show no signs of habituation.

## 4 Methods

### 4.1 Data pre-processing

The approach of analysis requires the MEG signal segmentation on *T* equal-length fragments, estimation of the spectral power for each segments separately and finally the statistical analysis of time-dependent properties of series of trial-segregated Fourier coefficients calculated for given frequencies. Similar, yet not equivalent algorithm was applied in studies presented in [39, 28].

To evaluate possible connection of trial-to-trial variability of MEG power spectrum with the measurement paradigm (tonal stimulation, stimulation by FM tones with or without categorization task, spontaneous activity) the registered signals were divided into segments time-locked to the stimulus (or to the beginning of the examination in the case of the measurement without stimuli) and separated from other segments by the same time interval being the difference between the fixed length of each trial (2 sec) and the length of the segments, like in [31] – see Fig. 2 a). Spectral power for consecutive trials of digital MEG signal was calculated as the discrete Fourier transform using the fast Fourier transform (FFT) algorithm (see for example [12]). Furthermore, a sequences of selected coefficients of these spectra *z_t_, t* = 1,…,*T*, were derived for some fixes frequency *f*. Note that only one spectral coefficient for each segment was selected at a time – see Fig. 2 b).

**Figure 2:**
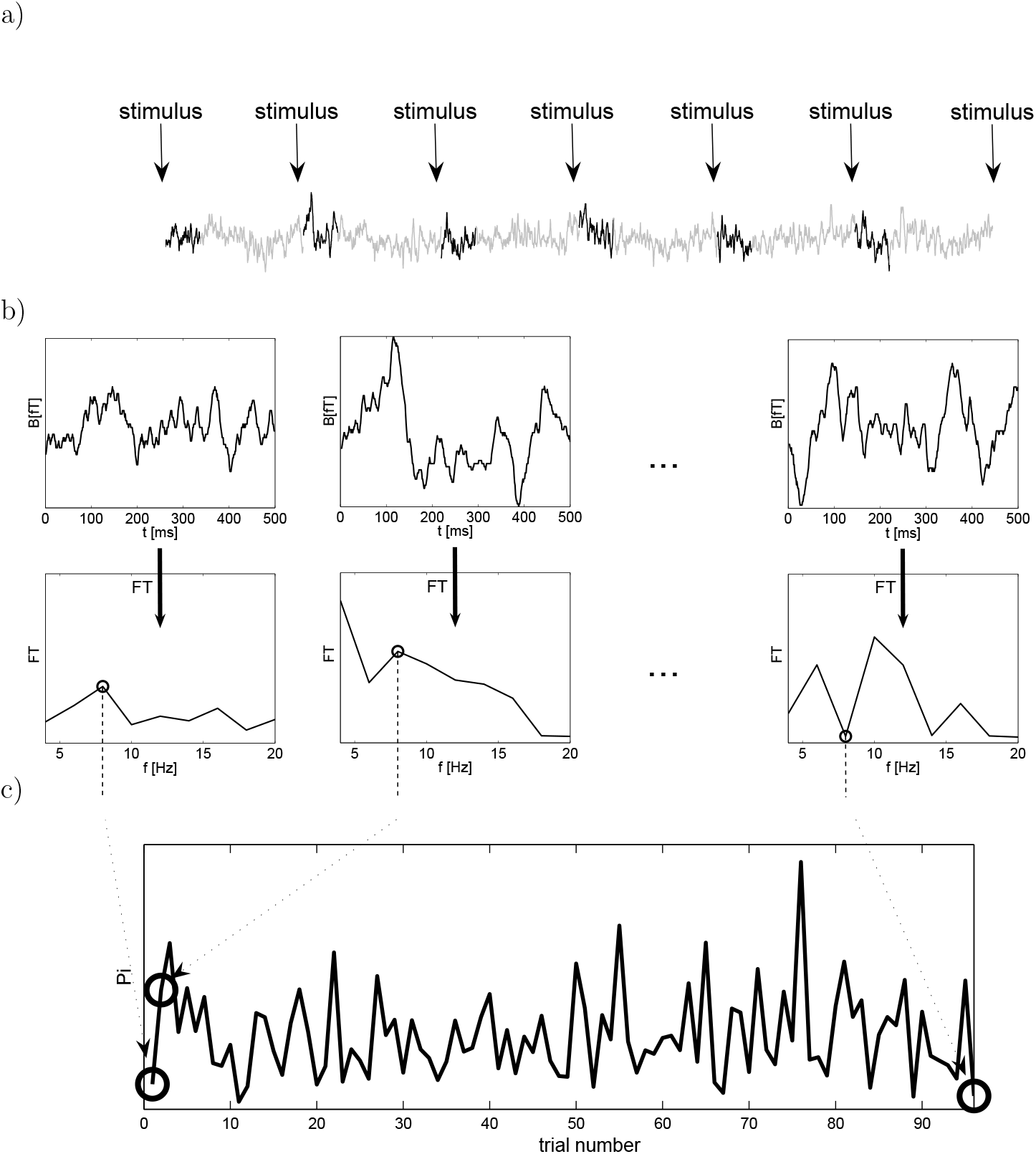
The scheme of computing the trial-to-trial variability time series of the MEG power spectrum.

Taking into account the frequency resolution and the nonstationary character of the data it was decided to perform analysis for two lengths of the aforementioned segments. Important was to use rather short segments in order to diminish the time error of the Fourier estimation and to maximize the signal-to-noise ratio of the Fourier transformed data (this is because the stimulus-related activity was assumed to last relatively shortly after the stimulus onset), yet, it was also tried to preserve a reasonable frequency resolution, which increases with increasing the length of the analyzed signal to be transformed. Therefore, the signals of both 250 ms and 500 ms duration were analyzed. The 250 ms signals contain the evoked components but not the waveforms related to endogenous (e.g., cognitive) activity. The 500 ms signals enable investigation of all event-related waveforms.

The time series analysis of a trial-to-trial power spectrum variability was performed for frequencies from the dominant range 8—12 Hz what is the alpha activity frequency band. It was obtained 148 (the number of MEG channels) series for each kind of stimulation and each of the selected signal durations. For the 500 ms signals the FFT spectra revealed three significant peaks, at 8 Hz, 10 Hz, and at 12 Hz, whereas the spectra of the 250 ms signals demonstrated two clear peaks — at 8 Hz and at 12 Hz. This is because of the inherent frequency resolution of the fast Fourier transform, which for these signals equaled 2 Hz (1/500ms^−1^) and 4 Hz (1/250ms^−1^), respectively.

Concluding, the obtained results are a sequence of the Fourier coeffcients for a fixed frequency calculated for each single-trial separately – see Fig. 2 c). Realizations for all trials form the stochastic process *Z_t_*.

The subsequent steps in Fig. 2 show the following procedures.

a. The original MEG signal segmentation according to stimulus onset (or equivalent interval of time): here a fragment of single-channel MEG registered during tonal stimulation is shown, the stimulus onsets are pointed by arrows and the marked segments (500 ms samples in this example) are selected to compute their spectra.
b. The single-trials MEG selected in step a) transformed with the use of FFT to the frequency domain; for a fixed frequency (8 Hz in this case) the time series of the consecutive Fourier coefficients is prepared.
c. The time series *z_t_* being the realization of the trial-to-trial-variability of the MEG power spectrum and subject to further analysis.

This calculations were performed for each of 148 MEG channel separately, and for all fixed frequencies from the dominant range 8—12 Hz (taking into account the spectral resolution) separately.

### 4.2 Mathematical model

For the stochastic process *Z_t_* let us denote its realizations by *z_t_*. If time t is discrete, *t* = 0, ±1,…, this process is the time series. Such process is weakly stationary if

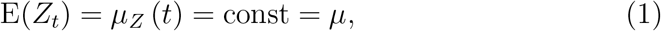

and

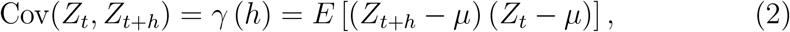

for *t* = 0, ±1,…, *h* = 0, ±1,…. In other words, it means that the meanvalue function is independent of time *t*, and that the covariance function depends on *r* and *s* only through their difference |*r* – *s*|, so it is independent of *t* for each *h* too.

The spectral density of a stationary process *Z_t_* specifies the frequency decomposition of the autocovariance function. Suppose that *Z_t_* is a zero-mean stationary time series with autocovariance function *γ*. If 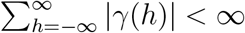 is satisfied, the process has the representation

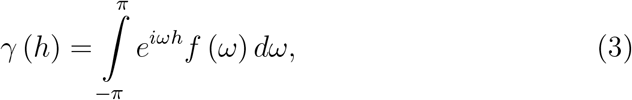

for *h* = 0, ±1, ±2,…, *e^iω^* = cos (*ω*) + *i* sin (*ω*), and 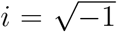. Equation (3) is the inverse transform of the spectral density of *Z_t_* defined as

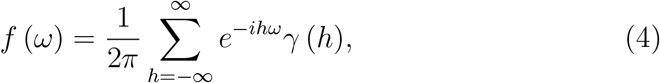

where –*π* ≤ *ω* ≤ *π*. The spectral density is an analogue to the probability density function; for all *ω* ∈ (–*π, π*], *f* (*ω*) ≥ 0 and it is even, i.e., *f* (*ω*) = *f*(–*ω*).

The relation between the spectral density function and the variance of the process {*Z_t_*} is

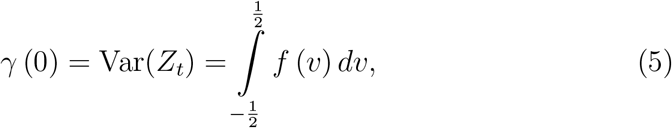

where *v* = *ω*/2*π* is the frequency and –1/2 ≤ *v* ≤ 1/2.

To transform one function into another that is the frequency-domain representation of the original function, we use the Fourier transforms (FTs). Discrete Fourier transform (DFT) is given by

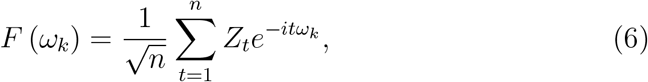

where *n* is the number of observations of a time series *Z_t_*, 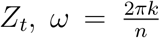, and 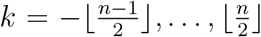, where ⌊*y*⌋ denotes the largest integer smaller than or equal to *y*. Then, each component of an observable time series *Z_t_* can be written as

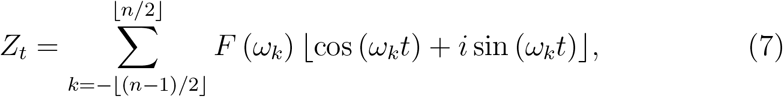

where *t* = 1,…, *n*, which shows that *Z_t_* can be represented as a linear combination of sine waves with frequencies *ω_k_*.

After procedure based on FFT algorithm, described in subsection 4.1, the obtained sequence of the selected coefficients of the single-trial power spectra for a fixed frequency will be now the time series *z_t_* being realization of the stochastic process *Z_t_* representing time-varying brain activity in frequency domain. Now, let us to present the procedure we propose to infer the *z_t_* properties and to fitting an adequate time series model to it. These procedure is given in a diagram form in Fig. 3.

**Figure 3:**
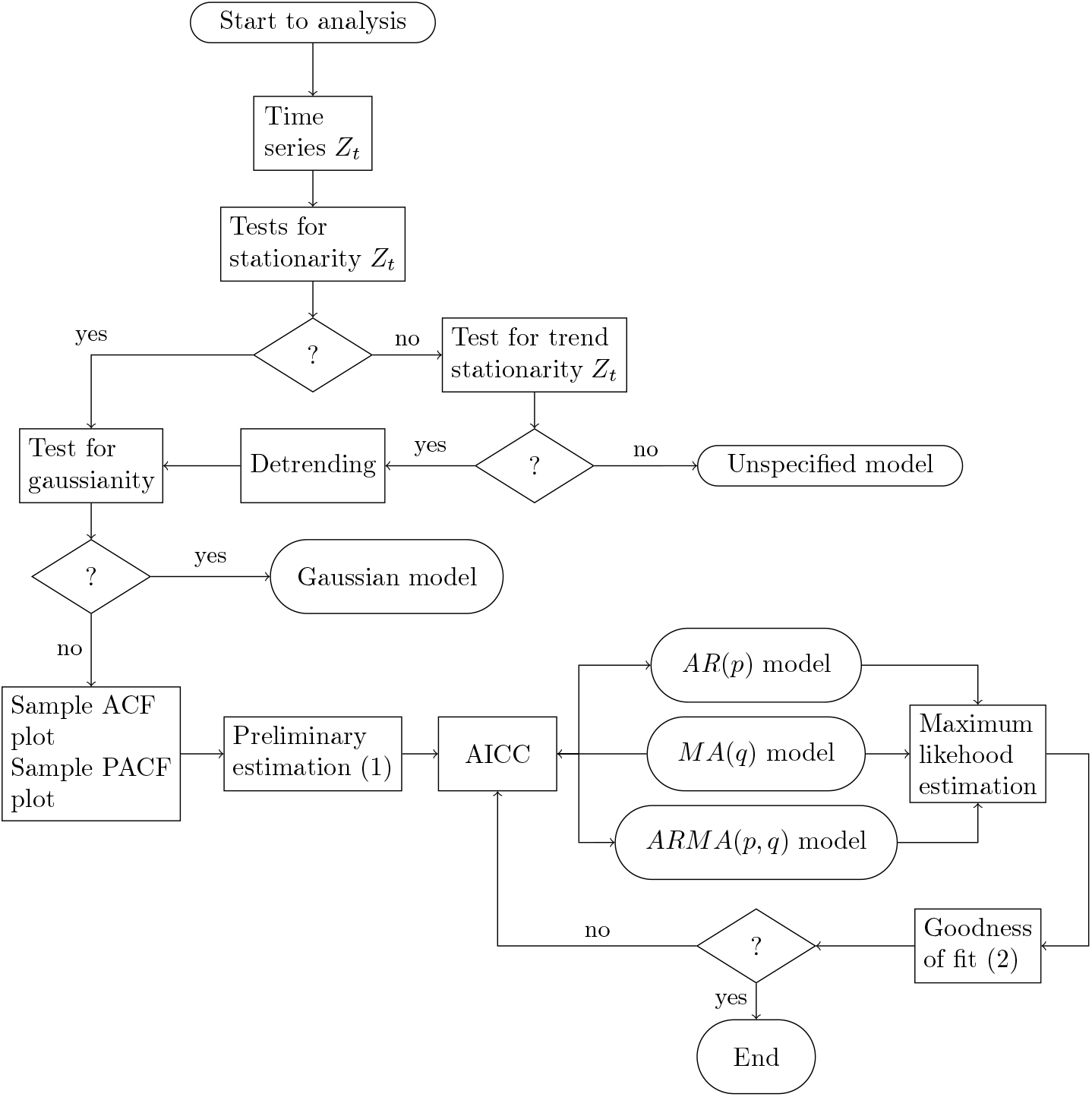
The statistical inference as the decision tree and the *z_t_* time series modeling procedure. Explanations in the text.

In order to recognize the stochastic nature of *Z_t_* it was crucial to verify their stationarity. In this case, similar to [31] were applied: the Kwiatkowski–Phillips–Schmidt–Shin (KPSS) test for level-stationarity and trend-stationarity [35], the Phillips–Perron (PP) test for unit root [43], and the White test for heteroscedasticity (variance-nonstationarity) [54]. We used these tests in the beginning of time series analysis, and if the trend-nonstationarity is detected, we made detrending to its elimination, as it is shown in a upper part of the decision tree from Fig. 3. Besides, the Jarque–Bera test [24] to verify the hypothesis about normality of the analysed series of spectral coefficients was also applied. Whenever Gaussianity was found, an appropriate model was fitted from the class of autoregressive moving-average (ARMA) models of the orders *p, q* ∈ [0,10], which was justified by the obtained form of the sample autocorrelation function (ACF) and the sample partial autocorrelation function (PACF) plots. Time series {*Z_t_*} is an (ARMA(*p, q*)) process:

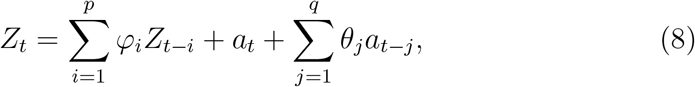

with some constant coefficients *φ_i_* and *θ_i_*, where {*a_t_*} is a white noise [10].

Fitting the time series model is a complex, iterative procedure and cannot be full described here, yet we introduce it shortly in a lower part of the diagram from Fig. 3. In this work the methods given in detail in handbook [11] were applied. For preliminary estimation (Fig. 3 (1)) of the parameters of ARMA models we applied: the Yule-Walker estimation and the Burg’s algorithm for pure autoregressive (AR) processes, the innovations algorithm and the Hannan-Rissanen algorithm for pure moving-average (MA) models, and the last-mentioned method for mixed (ARMA) models. All of these algorithms are described in [11]. Estimators of the model parameters were obtained by maximizing the Gaussian likelihood of the ARMA process. The major criterion for the selection of orders of the estimated model is minimization of the bias-corrected Akaike criterion (AICC) value, what is used in this studies too. At the end, we performed the diagnostic checking to verify the goodness of fit of a statistical model to a set of data (Fig. 3 (2)). We applied it by testing the residuals of the model, which should have the white noise properties if model is appropriate fitted. Therefore we applied varies tests for randomness (the Ljung-Box test, The McLeod–Li test, the turning points test, the difference-sign test, the rank test and the Jarque–Bera test) exactly as it is given in [11].

Finally, let us to remind that the procedure described above, leading to the adequate time series model of the trial-to-trial variability of the MEG power spectrum was performed for each experiment paradigm, each single MEG channel, each trial length and each of the fixed frequencies separately.

## 5 Results

### 5.1 Inference about stationarity

The stationarity of the series of Fourier coefficients were tested independently for all channels and measurement sessions. It verified the presumptions that the character of power spectrum trial-to-trial variability is related to the functional state of the brain. Such correlation should be observable in the predominant frequency component of the MEG signals. All the results of hypotheses testing are shown in Table 1.

**Table 1:** Results of testing the trial-to-trial variability of selected power spectra coefficients. Used are: the Kwiatkowski–Phillips–Schmidt–Schin (KPSS) test, the Phillips–Perron (PP) test, and the White test. Notation: LS — level-stationary, TS — trend-nonstationary, UR — unit root, VS — variance-stationary (homoscedastic), VNS — variance-nonstationary (heteroscedastic).

| experimental<br>paradigm | trial<br>duration<br>[ms] | frequency<br>[Hz] | % of rejection of statistical test |  |  |  |
| --- | --- | --- | --- | --- | --- | --- |
|  |  |  | KPSS |  | PP | White |
| | | | $H_0$: LS<br>$H_1$: UR | $H_0$: TS<br>$H_1$: UR | $H_0$: UR<br>$H_1$: LS | $H_0$: VS<br>$H_1$: VNS |
| no stimuli<br>(session 1) | 250 | 8 | 3.4 | 0 | 95.9 | 0 |
|  |  | 12 | 0.7 | 0.7 | 98.0 | 0.7 |
|  | 500 | 8 | 5.4 | 1.4 | 97.3 | 0 |
|  |  | 10 | 1.4 | 0 | 95.9 | 0.7 |
|  |  | 12 | 8.1 | 5.4 | 91.9 | 3.4 |
| 1 kHz sound<br>stimulation | 250 | 8 | 6.8 | 2.0 | 91.2 | 4.7 |
|  |  | 12 | 4.7 | 2.7 | 92.6 | 2.0 |
|  | 500 | 8 | 0 | 0 | 100 | 1.4 |
|  |  | 10 | 14.9 | 3.4 | 96.6 | 0 |
|  |  | 12 | 2.0 | 0 | 98.0 | 0.7 |
| no stimuli<br>(session 2) | 250 | 8 | 10.1 | 2.0 | 82.4 | 0.7 |
|  |  | 12 | 1.4 | 1.4 | 97.3 | 0 |
|  | 500 | 8 | 3.4 | 2.7 | 97.3 | 3.4 |
|  |  | 10 | 6.8 | 1.4 | 93.2 | 0 |
|  |  | 12 | 0 | 0 | 94.6 | 2.0 |
| FM/E<br>(exposure<br>condition) | 250 | 8 | 2.0 | 0.7 | 99.3 | 0 |
|  |  | 12 | 12.2 | 7.4 | 93.2 | 2.7 |
|  | 500 | 8 | 8.1 | 8.1 | 79.0 | 0 |
|  |  | 10 | 2.7 | 2.0 | 94.6 | 5.4 |
|  |  | 12 | 7.4 | 1.4 | 93.2 | 1.4 |
| FM/T<br>(task<br>condition) | 250 | 8 | 10.1 | 3.4 | 88.5 | 2.0 |
|  |  | 12 | 4.7 | 2.0 | 94.6 | 0 |
|  | 500 | 8 | 0.7 | 0 | 84.5 | 10.1 |
|  |  | 10 | 4.1 | 2.7 | 95.3 | 14.9 |
|  |  | 12 | 14.2 | 4.1 | 94.0 | 0 |

The obtained results show that the trial-to-trial variability of the power spectrum selected coefficients is evidently stationary in the wide sense. This property seems to be independent of the length of the analyzed signals, but the influence of the frequency resolution remains unknown. The highest percentage of rejecting the null hypothesis *H*_0_ was observed for the Phillips– Perron test, and the percentage of *H*_0_ acceptance is not fully comparable with the results obtained with the adequate version of the KPSS test (in 19 of 25 cases the percentage of accepting the PP-test hypothesis about the existence of a unit root is higher than the percentage of rejecting the KPSS-test hypothesis about trend-stationarity; there is only 1 opposite situation). The reason for this is the test power. The Kwiatkowski–Phillips–Schmidt–Schin test is a test of high power, and was proposed as an alternative for the unit-root tests due to the fact that those tests were known to fail to reject the null hypothesis of a unit root for many time series [35]. Therefore, authors believes that the results of the KPSS test are more objective. Having compared the results obtained with the two versions of the KPSS test the presence of trend-nonstationary in the analyzed data was considered only in a very small number of channels. The existence of the variance-nonstationarity was generally not proved either. Nor could be find any differences between the results of the two segment lengths. No significant differences between the results of for the power spectrum trial-to-trial variability at the selected frequencies were observed either. Finally, no clear channel-dependent properties were found.

### 5.2 ARMA modeling

The results for normality testing and time series modeling of the trial-to-trial variability of the MEG power spectrum are presented in Table 2.

**Table 2:**
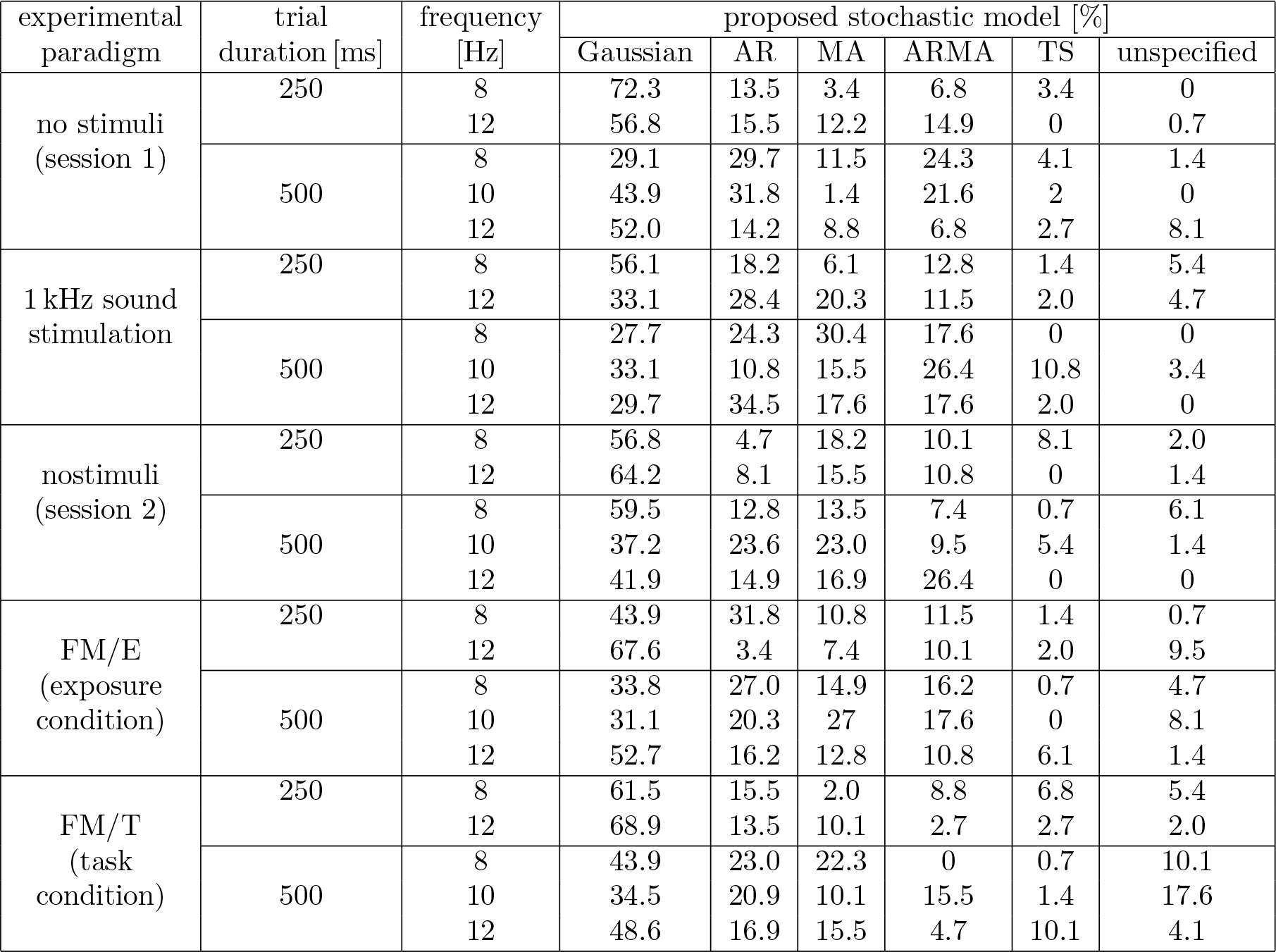
Results of testing for normality and of ARMA modeling of the trial-to-trial variability of the power spectrum. The percentage of revealing the normal distribution stems from the Jarque–Bera test. Notation as in Table 1.

It shows that the percentage of rejecting *H*_0_ in the Jarque–Bera test is a little smaller for the measurement without stimulation compared to the other experiments. It depends on the duration of the data segments taken for FFT, and longer signals usually have larger percentage of rejecting the hypothesis about Gaussianity. One exception is the power spectrum trial-to-trial variability of the MEG signals registered for the categorization-task paradigm. For those signals the averaged percentage of rejecting *H*_0_ by the Jarque–Bera test computed for all the analyzed frequencies (34.8% for the 250 ms and 57.7% for the 500 ms window) is similar to the results obtained for the ongoing activity (37.5% and 65.1%, respectively).

If not Gaussian, the ARMA model can be fitted in the vast majority of the analyzed time series of the power spectrum trial-to-trial variability. However, the percentage distribution of the proposed pure AR, pure MA, and mixed ARMA models is different for each of the examined experimental paradigms, and dependent on the length of the data segment. Moreover, the orders of the fitted models do not reveal any correlation with the type of stimulation. Nevertheless, it can be said that the trial-to-trial variability of the series of selected Fourier coefficients is generally characterized by low-order stochastic processes, because median of the model order was equal to: *p* = 3 for the AR, *q* = 2 for the MA, and *p* = 2, *q* = 1 for the mixed ARMA model or by a Gaussian process. It means that MEG spectral power in the given time window depends on only 2 or 3 previous observations. Additionally, the need to estimate only 2-3 coefficients proves the low level of complexity of the model and the simplicity of calculations.

A small percentage of the analyzed time series cannot be described by models from the ARMA class due to their nonstationarity. Fortunately, approximately half of them contain deterministic trend, therefore the trendplus-noise model can be proposed for their description. Finally less than 5% of all time series remained undescribed. It may be essential to note that the percentage of such “unknown” series increases relatively for the FM stimulation. It is especially visible for the 10 Hz frequency and distinct for the longer segments, which contain the cognitive event-related components. For these data this percentage reached the maximum of 17.6%.

Unfortunately, no correlation between the particular time series model characteristic and the MEG-channel localization over head was found.

## 6 Discussion

Although the physiological MEG signals recorded during the presented complex acoustic experiment are in principle nonstationary in time domain, the weakly stationary character of their power spectra exists at several frequencies. Our results suggest it is the common property for that kind of signals, rather independent from the functional state of brain i.e. recognition and classification of cognitive stimuli (event-related responses), processing of au-diosensory information (auditory evoked fields) or rest state (spontaneous MEG).

In some ways this observation is similar to the results obtained in [28] or [5], but presented here conclusions are somewhat different. The authors of the aforementioned articles claimed that their observations suggested that the brain signals are stationary, which is not really true [31]. In our opinion, short-time MEG signals may be nonstationary due to the time-varying spectrum, but the variability of their spectrum is the stochastic process, overwhelmingly with weakly stationary properties.

Applying the method of the signal transformation from the time domain to the frequency domain presented in this paper, the time-variability of the MEG spectrum mostly has a Gaussian distribution or could be described by the stationary ARMA time series models. The calculation of the parameters of these models turns out to be relatively simple and the obtained residua meet the criteria of a proper fit of the time series model to the data. In this way we obtained the effective models for MEG trial-to-trial spectral variability with medial 2—3 coefficients what shows that the current spectral power in a given frequency, calculated from 250ms or 500ms epoch lengths, depends on 2—3 previous epochs only. Taking into account the 2s trial length, the spectral power in a given trial depends on its value from the 4—6 seconds before.

Only a small percentage of the trial-to-trial power spectra changes in a different way, i.e. still presents nonstationarity. However, it should be emphasized that these cases concerned mainly longer fragments of MEG signals registered during the task condition with FM tones stimulation. This suggests that nonstationarity may be the result of a cognitive process and thus, in a way, be a marker of a significant state change in the neural network related to the classification of the direction of the frequency tone modulation. If so, the question remains why it is observed in such a small percentage of the data? Referring critically to the presented research, it is possible that the particular estimation method used for this studies is responsible for this. The temporal resolution of the Fourier transform is limited by the length of a time window to be transformed, as the FFT provides only one power spectrum for such a window. The frequency resolution is limited too, as the lowest frequency to be observed is the inverse of a time-window length. Therefore, the intrinsic limitations of the Fourier transform might have biased our results.

On the other hand, the presented algorithm of encephalographic data analysis is not computationally complicated and allows to obtain a low-complexity stochastic model of the EEG/MEG spectrum variability. Therefore, it could be useful in some theoretical neuroscience. Exemplary, when applied to the data presenting the habituation effect, we expect ARMA plus trend nonstationary models of their trial-to-trial spectra variability, which are also easy to estimate. It would also be worthwhile to investigate whether the parameters of the time-varying spectrum models will be different in particular pathologies. The proposed method of analysis may also find applications in the research where prediction of spectral changes plays important role. Usually the learning mechanism is used to estimate the required statistics of encephalographic data modeled by a complex network and a lot of research has been devoted to this issue [15, 40]. It should be interesting how the deep learning methods solve the problem of parametrization the trial-to-trial EEG/MEG signals variability – in the presented work this procedure required the researcher to make decisions in an iterative time series modeling algorithm (see the Fig. 3). Also, we see a lot of possible applications for it. The EEG characteristics prediction is helpful in versatile neurostimulation applications, such as the brain-machine interface based techniques [1, 13, 9] or other EEG-cooperating methods like electrical brain stimulation [36]. Methods of predicting changes in EEG, especially when applied online, can be of a great clinical utility. An appropriate model of spontaneous or evoked brain activity gives the possibility of forecasting the state-changes, what is crucial for patients in coma due to epileptic state, strokes or brain injuries [46]. So far, the best solutions in this area have been achieved in epileptic seizure prediction [55, 20] or prognostication in hypoxic-ischemic brain injury [2]. In this place let us to notice that the AR time series models are already used in forecasting of the alpha oscillatory phase of EEG signals [48] and the realtime epileptic seizure prediction [14]. We hope our research may support above mentioned and similar studies.

The proposed approach is by no means the final solution to the problem of describing the trial-to-trial variability of the EEG/MEG spectrum. We consider it is necessary to improve the methodology of such a time series modeling. Earlier we noted that the spectral density estimation method (FFT in our case) may bias the results. One can consider the use of wavelets or the matching pursuit (MP) algorithm for this purpose, as in the work [30], but then first of all approximations of the nonstationary spectrum are achieved and the ARMA model serves only to describe the residual noise. Another improvement would be to use multivariate state-space time series models instead of thier univariate version, similary like in works [18, 37]. Such models keeps the characteristics of the signals from all EEG leads or MEG channels at once, taking into account the cross-channel correlations.

## 7 Conclusions

The nonstationarity of MEG is important for understanding the brain functioning on each level of neuronal network. In this paper we investigated the very short MEG signals with the proven variance-nonstationary character and representing rest state in comparison to the auditory information processing at its few levels: from passive listening of simple tonal and frequency modulated stimuli to the single-trial brain responses related to their recognition and classification. By preparation the trial-to-trial spectral time series we investigated their time-variability at the dominant alpha (8—12 Hz) Fourier frequencies and we found the similarities independently of the functional state of brain. The weakly stationary character is not found in only a small percentage of the trial-to-trial power spectra time series, mostly related to the classification of the direction of the frequency tone modulation. This suggests that nonstationarity may be the result of a cognitive process underlies the generation of an auditory evoked response.

We conclude there could be the information about neural processing of event-related information that exist in the spectrum variability and it is possible to parametrise it by rather simple stochastic models. Finally we proposed univariate ARMA class stationary time series models of a low-order for their description. Thus, we showed that the spectral density of analyzed signals depends on only 2-3 previous trials.

We hope our basic research is interesting for the scientists studying prediction models of EEG/MEG changes. Of course we are focused on spectral properties not the forecasting or reconstruction the encephalographic waveforms in time domain.

We believe that the presented method may prove useful in some theoretical research and increase knowledge of the properties of brain signals, which may be of benefit to more advanced research in neuroscience, which has been discussed. From a clinical point of view, this work could contribute to the development of a new method of predicting changes in EEG spectrum in epilepsy or neuromonitoring patients.

## Acknowledgements

Authors would like to thank colleagues from Leibniz Institute for Neurobiology in Magdeburg (Germany), Reinchard König and Cezary Sielużycki for providing the data for analysis and Norman Zacharias for the realization of the MEG measurement.

